# Isobaric Tagging and Data Independent Acquisition as Complementary Strategies for Proteome Profiling on an Orbitrap Astral Mass Spectrometer

**DOI:** 10.1101/2024.12.17.628765

**Authors:** Xinyue Liu, Shane L. Dawson, Steven P. Gygi, Joao A. Paulo

## Abstract

Comprehensive global proteome profiling that is amenable to high throughput processing will broaden our understanding of complex biological systems. Here, we evaluated two leading mass spectrometry techniques, Data Independent Acquisition (DIA) and Tandem Mass Tagging (TMT), for extensive protein abundance profiling. DIA provides label-free quantification with a broad dynamic range, while TMT enables multiplexed analysis using isobaric tags for efficient cross-sample comparisons. We analyzed 18 samples, including four cell lines (IHCF, HCT116, HeLa, MCF7) under standard growth conditions, in addition to IHCF treated with two H₂O₂ concentrations, all in triplicate. Experiments were conducted on an Orbitrap Astral mass spectrometer, employing Field Asymmetric Ion Mobility Spectrometry (FAIMS). Despite utilizing different acquisition strategies, both the DIA and TMT approaches achieved comparable proteome depth and quantitative consistency, with each method quantifying over 10,000 proteins across all samples, with slightly more protein-level precision for the TMT strategy. Relative abundance correlation analysis showed strong agreement at both peptide and protein levels. Our findings highlight the complementary strengths of DIA and TMT for high-coverage proteomic studies, providing flexibility in method selection based on specific experimental needs.

## INTRODUCTION

Accurate quantification of the global proteome is imperative for understanding complex biological systems. As proteomics studies expand in scale and complexity, high-throughput capabilities are essential to minimize time demands for large-scale analyses. Achieving deep proteome coverage, which includes detecting low-abundance proteins that are often vital to cellular functions, is equally important for comprehensive insights. Here, we highlight the complementary strengths of two prominent quantitative mass spectrometry techniques, Data Independent Acquisition (DIA) and Tandem Mass Tagging (TMT), for achieving accurate, high-coverage protein profiling.

DIA-based workflows fragment all precursor ions within a specified mass-to-charge (m/z) range, capturing comprehensive data that enables broad dynamic range quantification across samples [1]. This label-free acquisition strategy minimizes sample processing requirements while allowing flexible, retrospective data analysis [2]. DIA is particularly advantageous for reproducibly quantifying low-abundance peptides with high reliability [3], making it ideal for studies requiring consistent, comprehensive proteome coverage.

Alternatively, isobaric tagging strategies, such as TMT [4], iTRAQ [5], and DiLeu [6, 7], facilitate sample multiplexing, enabling higher throughput and minimizing missing values. These methods label peptides with isobaric tags that release quantifiable reporter ions upon fragmentation, allowing the simultaneous analysis of multiple conditions in a single experiment. The high multiplexing capability of TMT (currently up to 35 samples [8]) is advantageous for complex sample comparisons. However, reliance on reporter ions can introduce noise and interference, which may limit the quantification of low-abundance peptides. Specifically, ion interference in TMT can lead to ratio compression, distorting relative protein abundance [9, 10].

Despite these methodological differences, both TMT and DIA significantly advance proteomics by offering distinct benefits, with some challenges. A general understanding of these techniques’ strengths and limitations aids in designing robust proteomic experiments that yield valuable insights. Here, we applied both approaches to 18 samples, which included four distinct cell lines: IHCF, HCT116, HeLa, and MCF7. Additionally, the IHCF cell line was treated with two different concentrations of H₂O₂. Biological triplicates were prepared for each condition, and each sample was divided at the peptide level to enable comparative analysis by both DIA and TMT strategies.

Our experiments were conducted on the Orbitrap Astral mass spectrometer, which integrates high-resolution mass analysis with high sensitivity, facilitating precise quantification of thousands of proteins from complex mixtures. For TMT samples, we employed Field Asymmetric Ion Mobility Spectrometry (FAIMS), which effectively reduces spectral complexity and enhances detection of low-abundance ions, a notable advantage in isobaric tag-based analyses. FAIMS is less commonly used with DIA, due in part to the potential reduction in data points per peak, but it remains feasible through modest adjustments to the instrument setup. Here, we further assessed whether DIA with a single FAIMS compensation voltage (CV) yielded comparable results to traditional DIA without FAIMS. Our bioinformatics analyses demonstrated that, despite differences in sample preparation and acquisition strategies, both DIA and TMT approaches provided similar depth and exhibited quantitative consistency within this experimental setup.

## MATERIALS AND METHODS

### Materials

Tandem mass tag (TMTpro) isobaric reagents were from ThermoFisher Scientific (Waltham, MA). Dulbecco’s modified Eagle’s medium (DMEM), both supplemented with 10% fetal bovine serum (FBS), were from LifeTechnologies (Waltham, MA). Trypsin was purchased from Pierce Biotechnology (Rockford, IL) and LysC from Wako Chemicals (Richmond, VA). Unless otherwise noted, all other chemicals were from Pierce Biotechnology (Rockford, IL). The Immortalized Human Corneal Fibroblast (IHCF, Cat. T0578) and the PrimGrow III medium (Cat. TM003) were purchased from Applied Biological Material Inc. (Richmond, BC, Canada). The remaining cell lines were purchased from ATCC (Manassas, VA).

### Cell growth and harvesting

Methods of cell growth and propagation for the IHCF cell line followed instructions provided by Applied Biological Materials Inc. In brief, the cell lines were propagated in PrimGrow III supplemented with 10% FBS. For the cells to be tested under oxidative stress, hydrogen peroxide was added at 40% cell confluency at 20 μM and 40 μM when refreshing the medium. Two days after treatment, the growth media was aspirated when the cells reached ∼90% confluency and the cells were washed thrice with ice-cold phosphate-buffered saline (PBS) before being dislodged with 0.25% Trypsin-EDTA. After neutralization using a complete growth medium, all cells were pelleted by centrifugation at 1,000 x *g* for 5 min at 4 °C and washed with ice-cold PBS twice.

Methods of cell growth and propagation for the remaining cell lines followed techniques utilized previously [11–13]. Briefly, these cell lines were propagated in DMEM supplemented with 10% FBS. The growth media was aspirated upon achieving ∼90% confluency. The cells were then washed thrice with ice-cold phosphate-buffered saline (PBS), dislodged with a non-enzymatic reagent, and harvested by trituration. After washing, all cells were pelleted by centrifugation at 1,000 x *g* for 5 min at 4 °C, and the supernatant was removed. The biological replicates used herein refer to cells harvested from different culture plates that have been propagated from a single frozen cell stock.

### Cell lysis and protein digestion

Three hundred microliters of native lysis buffer (PBS with 0.1% NP-40, pH 7.4) were added directly to each pellet for lysis. Cells were homogenized by 12 passes through a 21-gauge (1.25 inches long) needle. The homogenate was sedimented by centrifugation at 21,000 x *g* for 5 min and the supernatant was transferred to a new tube. Protein concentrations were determined using the bicinchoninic acid (BCA) assay (ThermoFisher Scientific). Proteins were subjected to disulfide bond reduction with 5 mM tris (2-carboxyethyl) phosphine (room temperature, 15 min) and alkylation with 10 mM iodoacetamide (room temperature, 20 min in the dark). Excess iodoacetamide was quenched with 10 mM dithiothreitol (room temperature, 15 min in the dark). Methanol-chloroform precipitation was performed prior to protease digestion [14]. Samples were resuspended in 200 mM EPPS, pH 8.5 and digested at room temperature for 14 h with LysC protease at a 100:1 protein-to-protease ratio. Trypsin was then added at a 100:1 protein-to-protease ratio and the reaction was incubated for 6 h at 37 °C.

### Tandem mass tag labeling (for TMT-based data acquisition)

TMTpro reagents (0.8 mg) were dissolved in anhydrous acetonitrile (40 μL), of which 6 μL was added to the peptides (25 µg) with 7 μL of acetonitrile to achieve a final concentration of approximately 30% (v/v). Following incubation at room temperature for 1 h, the reaction was quenched with hydroxylamine to a final concentration of 0.3% (v/v). TMTpro-labeled samples were pooled at a 1:1 ratio across all 18 samples. For each experiment, ∼400 µg of the pooled sample was vacuum centrifuged to near dryness and subjected to a C18 solid-phase extraction (SPE) column with a capacity of 100 mg (Sep-Pak, Waters).

### Off-line basic pH reversed-phase (BPRP) fractionation

We fractionated the pooled, labeled peptide sample using BPRP HPLC [15] on an Agilent 1260 pump equipped with a degasser and a UV detector (set at 220 and 280 nm wavelength). Peptides were subjected to a 50-min linear gradient from 5% to 35% acetonitrile in 10 mM ammonium bicarbonate pH 8 at a flow rate of 0.25 mL/min over an Agilent ZORBAX 300Extend C18 column (3.5 μm particles, 2.1 mm ID and 250 mm in length). The peptide mixture was fractionated into a total of 96 fractions, which were consolidated into 24 “super-fractions” [16]. Each super-fraction consisted of 4 fractions from the 96 well plate, corresponding to every 24^th^ fraction. Samples were subsequently acidified with 1% formic acid and vacuum centrifuged to near dryness. Each super-fraction was desalted via StageTip, dried again via vacuum centrifugation, and reconstituted in 5% acetonitrile and 5% formic acid for LC-FAIMS-MS^2^ processing.

### Mass spectrometric data collection (for TMT-based data acquisition)

Mass spectrometry data were collected using an Orbitrap Astral mass spectrometer (Thermo Fisher Scientific, San Jose, CA) coupled to a Neo Vanquish liquid chromatograph. Peptides (∼500 ng) from each super fraction were separated on a 110 cm µPAC C18 column (ThermoFisher Scientific). For each analysis, we loaded ∼0.5 μg. Peptides were separated using a 75-min method (60 min linear gradient) of 11 to 28% acetonitrile in 0.125% formic acid at a flow rate of 350 nL/min.

The scan sequence began with an Orbitrap MS^1^ spectrum with the following parameters: resolution 60,000, scan range 350−1350 Th, automatic gain control (AGC) target 200%, maximum injection time 50 ms, RF lens setting 50%, and centroid spectrum data type. FAIMS was enabled with compensation voltages (CVs): −35V, −45V, −55V, −60V, and −70V. We selected the top 35 precursors for MS^2^ analysis which consisted of HCD high-energy collision dissociation with the following parameters: Astral data acquisition (TMT on), AGC 100%, maximum injection time 25 ms, isolation window 0.4 Th, normalized collision energy (NCE) 35%, and centroid spectrum data type. In addition, unassigned and singly charged species and those >5+ were excluded from MS^2^ analysis and the dynamic exclusion was set to 15 s.

### Mass spectrometric data analysis (for TMT-based data acquisition)

Mass spectra were processed using a Comet-based in-house software pipeline. Database searching included all entries from the human UniProt database, which were concatenated with a reverse database composed of all protein sequences in the reversed order. Searches were performed using a 50- ppm precursor ion tolerance for total protein level profiling. The product ion tolerance was set to0.02 Da. These wide mass tolerance windows were selected to maximize sensitivity in conjunction with Comet searches and linear discriminant analysis (LDA) [17, 18]. TMTpro labels on lysine residues and peptide N-termini (+304.207 Da), as well as carbamidomethylation of cysteine residues (+57.021 Da), were set as static modifications, while oxidation of methionine residues (+15.995 Da) was set as a variable modification. Peptide-spectrum matches (PSMs) were adjusted to a 1% false discovery rate (FDR) [19, 20]. PSM filtering was performed using LDA [18] and then assembled further to a final protein-level FDR of 1% [20]. PSMs with a signal-to-noise value <1000, isolation purity <50%, and/or resolving power <40,000 for reporter ions were omitted from further analysis. Proteins were quantified by summing reporter ion counts across all matching PSMs, also as described previously [21]. Reporter ion intensities were adjusted to correct for the isotopic impurities of the different TMTpro reagents according to manufacturer specifications. The signal-to-noise (S/N) measurements of peptides assigned to each protein were summed and these values were normalized so that the sum of the signal for all proteins in each channel was equivalent to adjust for unequal protein loading. Finally, each protein abundance measurement was scaled such that the summed signal-to-noise for that protein across all channels equaled 100, thereby generating a relative abundance (RA) measurement. Data analysis and visualization were performed in Microsoft Excel or R.

### Mass spectrometric data collection (for DIA-based data acquisition)

Mass spectrometry data for the DIA experiments were also collected using an Orbitrap Astral mass spectrometer (Thermo Fisher Scientific, San Jose, CA) coupled with Neo Vanquish liquid chromatograph. Peptides (∼500 ng) from each super-fraction were separated on a 110 cm µ PAC C18 column (ThermoFisher Scientific) with a 75 min method (60 min linear gradient) of 11 to 28% acetonitrile in 0.125% formic acid with a flow rate of 350 nL/min.

The scan sequence began with an Orbitrap MS^1^ spectrum with the following parameters: resolution 240,000, scan range 380-980 Th, automatic gain control (AGC) target 500%, maximum injection time 50 ms, RF lens setting 50%, and centroid spectrum data type. FAIMS was enabled with a compensation voltage (CV) of −35V (“CV-35” dataset), or the FAIMS source was removed physically from the system (“NoFAIMS” dataset). Advanced peak determination (APD) was activated.

For DIA scans, the precursor mass range was 380-980 m/z. The DIA window was set to “auto” and the width to 2 m/z, which resulted in a duty cycle of 300 scan events. The collision energy was set to 27%. When FAIMS was used, the FAIMS voltage was set to −35V. The normalized AGC was set to 500% while the maximum injection time was set to 3 ms.

### Mass spectrometric data analysis (for DIA-based data acquisition)

Raw files were analyzed with Spectronaut 19.1.240806.62635 using the directDIA analysis type. The settings for Pulsar and the directDIA+ workflow were as follows: Trypsin/P and LysC as specific enzymes; peptide length from 7 to 52; max missed cleavages 2; Carbamidomethyl on cysteine as fixed modification and oxidation on methionine as a variable modification; FDRs at PSM, peptide and protein level all set to 0.01; minimum fragment relative intensity 1%; 3–6 fragments kept for each precursor. Data analysis and visualization were performed in Microsoft Excel or R.

## RESULTS and DISCUSSION

### Data were collected using DIA (with and without FAIMS) and TMT-based strategies

We applied Data-Independent Acquisition (DIA) and Tandem Mass Tag (TMT)-based strategies to 18 samples across four distinct cell lines (IHCF, HCT116, HeLa, and MCF7), with additional treatment of the IHCF line using two concentrations of H₂O₂, all in biological triplicate (**Fig. 1**). Each sample was lysed, reduced, alkylated, precipitated, and digested with LysC and trypsin prior to being divided for analysis by DIA and TMT-labeling according to the SL-TMT workflow [14].

**Figure 1:**
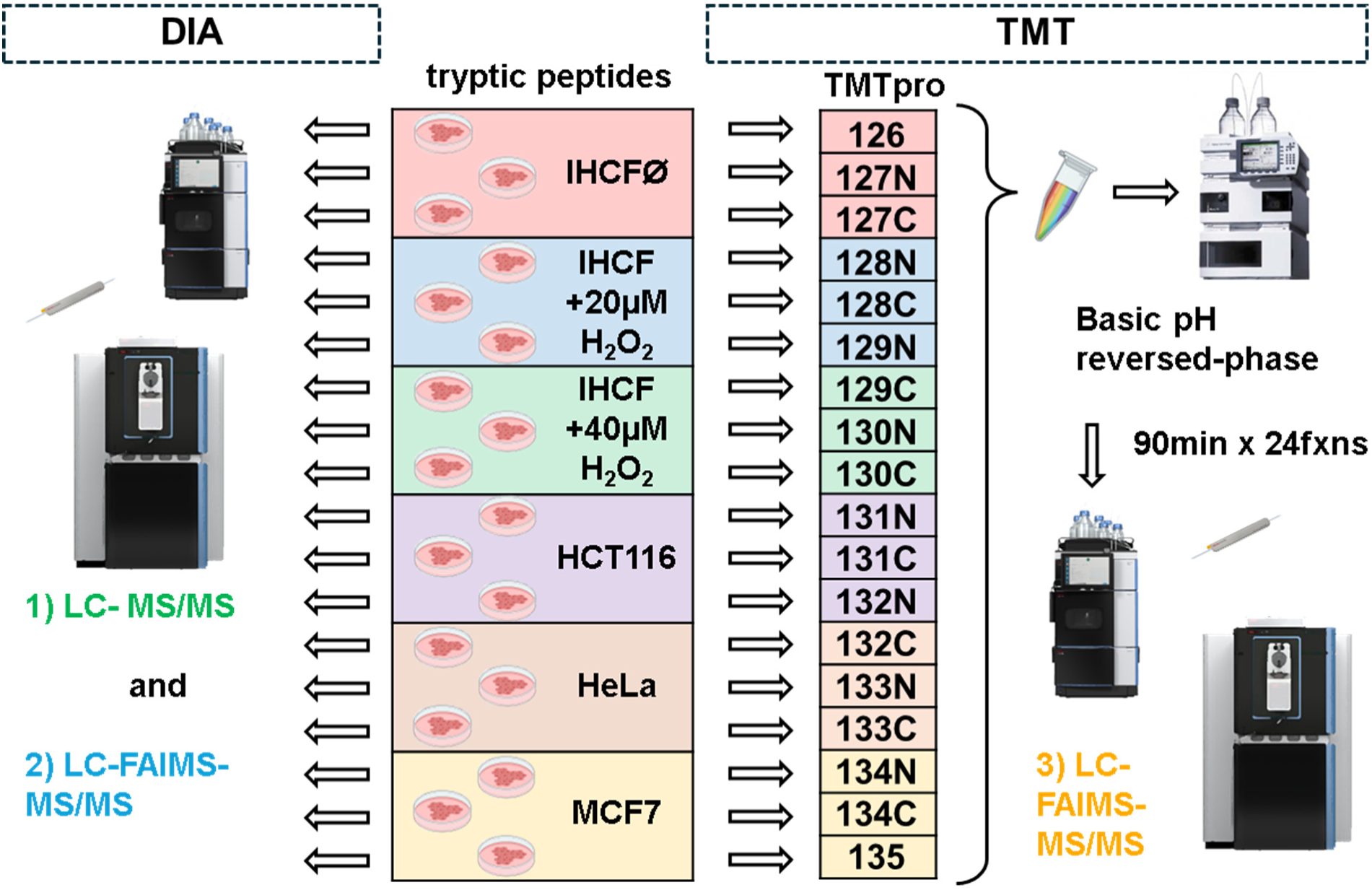
Workflow overview for processing 18 samples by DIA and TMT. Tryptic peptides were prepared from proteins extracted from reduced and alkylated cell lysates. Peptide pools were then aliquoted with 0.5 µg of each sample to be analyzed by DIA once without FAIMS and again with FAIMS (single CV of −35V). Aliquots (25 µg) from the same sample were labeled with TMTpro, pooled, and pre-fractionated with basic pH reversed-phase chromatography. The subsequent 24 concatenated super-fractions were analyzed by LC-FAIMS-MS/MS.

We collected three datasets using an Orbitrap Astral mass spectrometer with a micropillar array chromatography (µPAC) column. The two DIA datasets analyzed all 18 samples across 60-min gradients. We collected one DIA dataset without FAIMS, as is common practice, and acquired a second DIA dataset with FAIMS, applying a single compensation voltage (CV) setting of −35V. This CV was empirically determined to result in the highest number of quantified proteins for unlabeled samples. We noted that optimal CVs can vary between systems and recommend testing a range of CVs locally to achieve maximum data quality and depth. In addition, we acquired a third (TMT-based) dataset consisting of a fully fractionated 12-fraction TMTpro18-plex experiment. Data analysis for the DIA datasets was conducted using Spectronaut 19 with the directDIA analysis type, while the TMT-based data analysis was conducted using a Comet-based workflow.

### Proteome profiling with DIA with FAIMS quantified a similar number of proteins with highly correlated quantitative profiles compared to DIA without FAIMS

We sought to evaluate DIA on an Astral mass spectrometer both using conventional DIA (without FAIMS) and using a single FAIMS compensation voltage (CV). For TMT-based analyses, the use of FAIMS significantly decreases spectral complexity and therefore ion interference by gas-phase fractionation, leading to more accurate peptide and thus, protein quantification. Moreover, we have observed previously for TMT-based analyses that FAIMS increases the number of proteins, yet its inherent selectivity actually decreases the number of quantified peptides [22, 23].

We first compared the number of peptide-spectrum matches (PSMs) in each of the 18 samples (**Fig. 2A**). On average, over 150,000 PSMs were quantified in each sample of the NoFAIMS dataset, while ∼30% fewer, 100,000 PSMs were quantified in the CV-35 dataset. We used a Venn diagram to compare the overlap of total peptides between the two datasets (**Fig. 2B**), which revealed that only 48% of the quantified peptides overlapped. We also showed that >100,000 more (unique) peptides were quantified in the NoFAIMS dataset, while only ∼17,000 peptides were unique to the CV-35 dataset. This loss of peptides was expected as only 1 CV was used, and as such only precursors with ion mobility compatible with this CV (*i.e.,* FAIMS separates ions based on size, shape, charge, and other ion-specific properties) were analyzed [24, 25]. However, the difference between the two datasets nearly dissipated at the protein level (**Fig. 2C**). Here ∼10,000 proteins were quantified in each sample, with marginal differences between corresponding NoFAIMS and CV-35 analyzed samples. Likewise, the overlap at the protein level was noticeably greater than at the peptide level (**Fig. 2D**). Specifically, over 9,000 proteins (>67%) overlapped between the two data acquisition methods, with ∼18% and ∼15% of proteins that were unique to the NoFAIMS and CV-35 datasets, respectively. Overall, the number of proteins quantified were similar, even though the number of peptides was ∼30% lower in the NoFAIMS DIA dataset.

**Figure 2:**
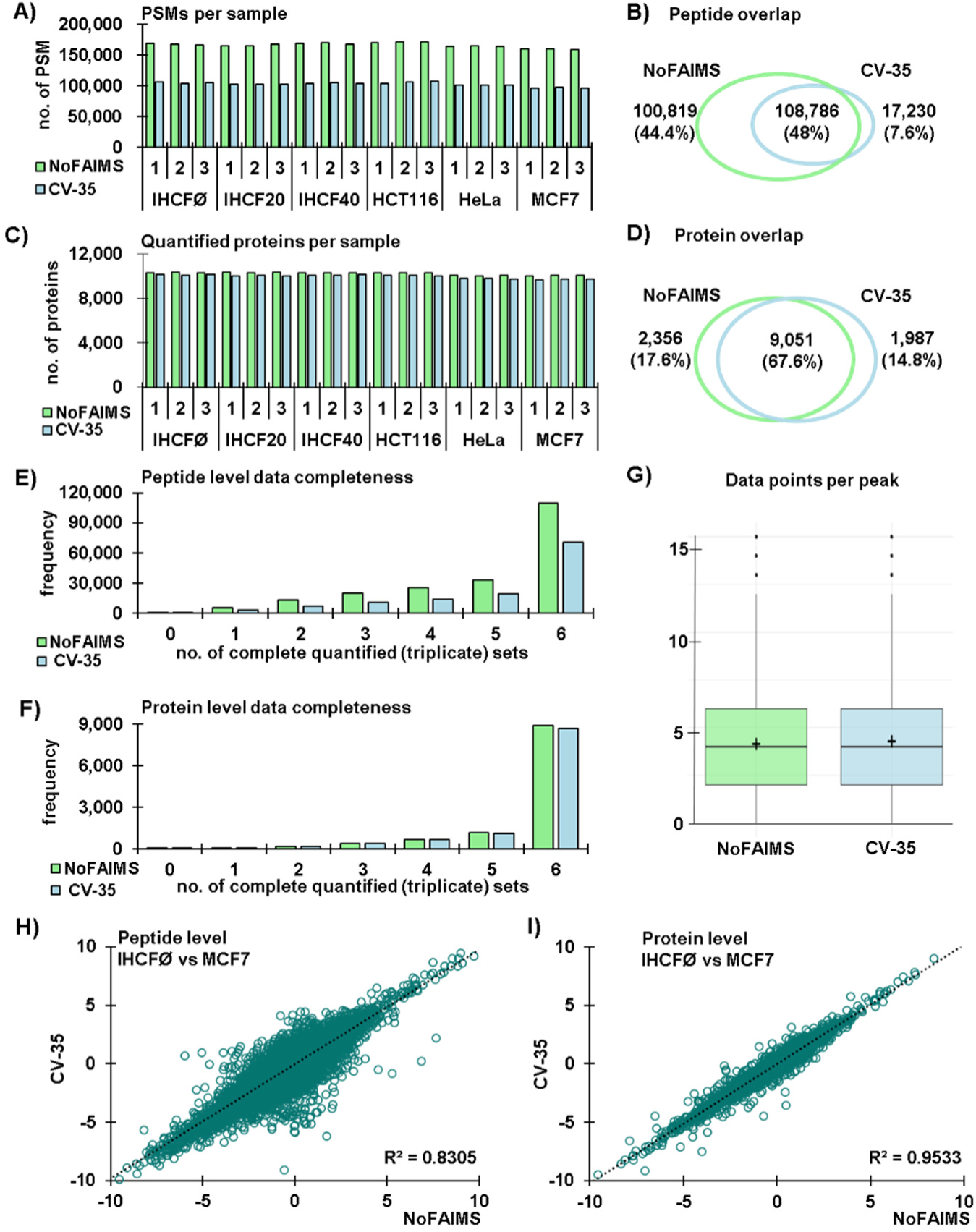
Comparison of DIA datasets acquired with no FAIMS and with one FAIMS compensation voltage (CV=-35V). **A)** Count of the total peptides quantified in each sample for data acquired without (“NoFAIMS”) and with FAIMS (“CV-35”). **B)** Venn diagram depicting the overlap in peptide sequences quantified by both DIA acquisition methods. **C)** Count of the proteins quantified in each sample for each DIA acquisition method. **D)** Venn diagram illustrating the protein-level overlap between DIA acquisition methods. **E)** Peptide level completeness for each DIA acquisition method. “Completeness” requires a measurement for each replicate for a given peptide. If no measurement is made for an entire set of triplicate samples, it does not count against completion, as it is assumed that the given protein is not expressed in that cell line. **F)** Protein level completeness for each DIA acquisition method using the same criteria as in (**E**). **G)** Box-and-whiskers plot depicting the number of data points for a given chromatographic peak. **H)** Correlation plot of the average fold change in abundance between IHCF and MCF7 cell lines for the two DIA acquisition methods at the peptide level. **I)** Corresponding correlation plot of the average fold change in abundance between IHCF and MCF7 cell lines at the protein level.

In addition to the depth of the analysis, data completion was also imperative as we aim to quantify proteins across all samples. In the case of this sample set, we investigated several very different cell lines and so we expected that not all proteins will be expressed in all cell lines. It followed that we defined data completion with respect to replicates, such that if no measurement was made for a set of triplicate samples, it did not count against completion. In total, the dataset consisted of six sets of triplicates. We noted that at the peptide level, most peptides were quantified in all six replicates for both the NoFAIMS and CV-35 datasets. The proteins in a given number of replicate sets steadily decreased as the number of replicate sets decreased (**Fig. 2E**). The same trend was seen at the protein level, with very little difference between the two DIA strategies (**Fig. 2F**). We can also define data completion more conservatively by the number of samples (out of the total 18) in which a given peptide was quantified in the DIA dataset. (**Fig. S1A**). We noted that for both the NoFAIMS and CV-35 datasets, most peptides were quantified in all 18 samples, with decreasing numbers quantified as the number of samples with a given peptide decreased. This result was reflected at the protein level, with an even greater percentage of proteins quantified in all 18 samples (**Fig. S1B**). Together with data completion, the number of data points per chromatographic peak was an important metric for data quality. We observed that despite fewer quantified peptides, both DIA strategies averaged approximately 4 data points per peak (**Fig. 2G**). In all, the data acquired by both DIA strategies were qualitatively similar.

We also assessed the quantitative correlation between the two DIA strategies. First, we compared the quantitative abundance differences at the peptide level. As an example, we compared the abundance changes for the IHCF and MCF7 cell lines, which we illustrated with a correlation plot (**Fig. 2H**). The Pearson R^2^ for this peptide-level comparison was 0.8305, signifying a high degree of correlation at the peptide level (**Fig. 2I**). It followed that at the protein level, the correlation was even higher at 0.9533. Overall, these data showed that the use of FAIMS with DIA data acquisition did not severely affect the number or quantitative consistency at the protein level, despite fewer quantified peptides.

### TMT-based data acquisition resulted in a similar number of quantified proteins with well-correlated quantitative profiles

Next, we compared our DIA datasets with a TMTpro18-plex dataset consisting of the same 18 peptide samples, which were now labeled with TMTpro reagents, pooled, and fractionated. This dataset was analyzed with FAIMS, as was recommended for MS2-based isobaric tag quantification to reduce interference [22, 23]. We first compared the proteins quantified by each data acquisition strategy (**Fig. 3A**). Over 11,000 proteins were quantified using the DIA methods, with nearly 9,000 proteins quantified in all replicates using the completeness criteria from **Fig. 2F**. These values were comparable to the 10,325 proteins quantified across all samples in the TMTpro18-plex experiment.

**Figure 3:**
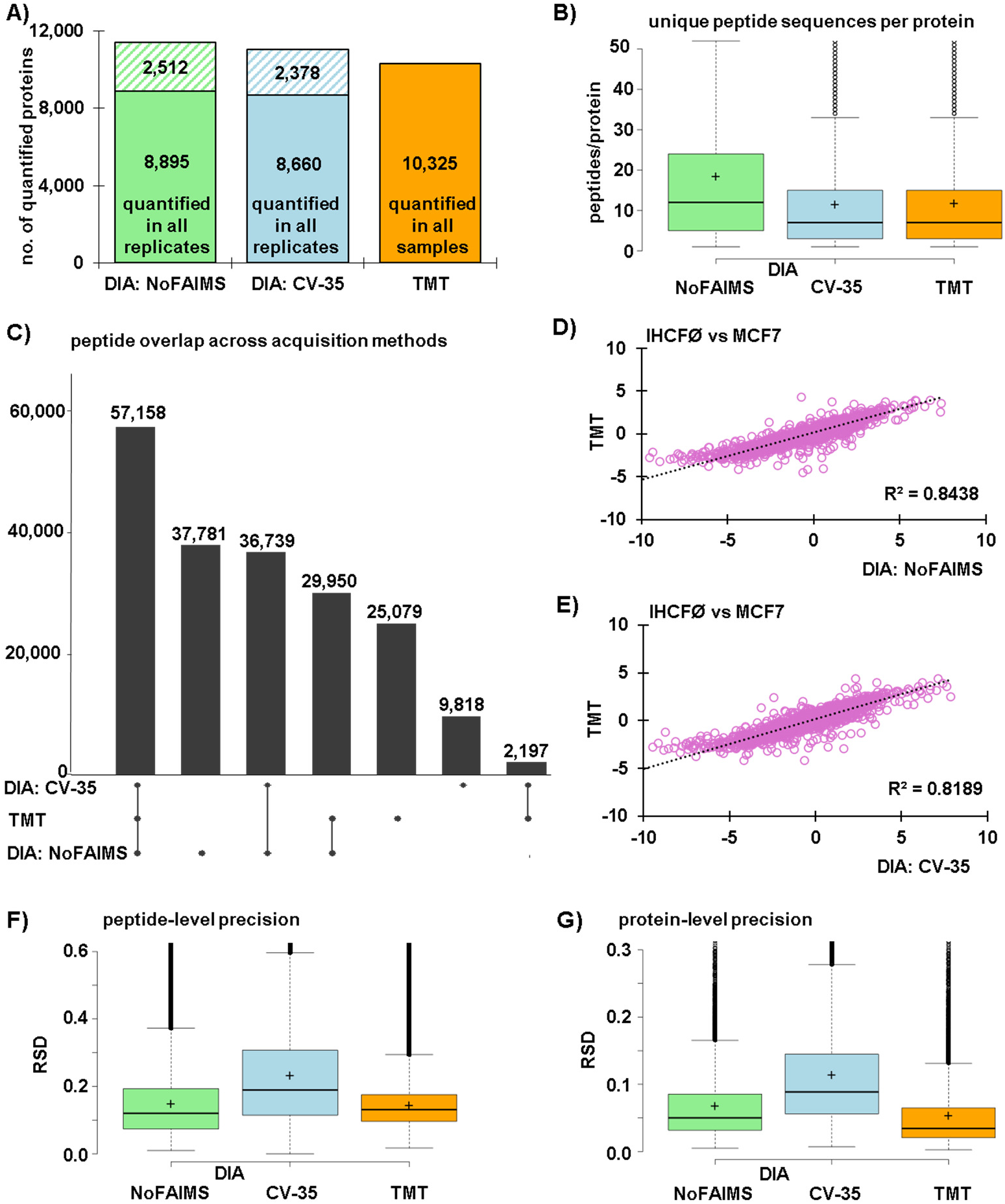
Comparison of DIA and TMT datasets acquired on an Orbitrap Astral mass spectrometer. **A)** Bar graph depicting the number of quantified proteins in all three datasets. The stacked bars for the DIA datasets include proteins in which all triplicates of a given condition had measurements (or all had no measurements) for a given protein (bottom portion, solid) and those with missing values (top portion, slashes). **B)** Box-and-whiskers plot illustrating the number of peptides with unique sequences quantified per protein. **C)** An UpSet plot showing the peptide overlap across the three data acquisition methods. Correlation plot of the fold change in abundance between IHCF and MCF7 cell lines at the peptide level for **D)** the TMT and DIA: NoFAIMS datasets, and **E)** the TMT and DIA: CV-35 datasets. Box-and-whiskers plot illustrating the distribution of the coefficient of variation (relative standard deviation, RSD) for triplicate measurements of each condition (or cell line) at the **F)** peptide and **G)** protein level for all three datasets.

We then examined the number of peptides per protein for these datasets (**Fig. 3B**). Proteins quantified in the DIA: NoFAIMS dataset were represented by an average of nearly 20 unique peptides. However, the two methods using FAIMS showed a similar distribution of unique peptides per protein, both averaging ∼10, highlighting further the equivalence of these two methods. In addition, we generated an UpSet plot to illustrate peptide overlap among the three data acquisition strategies (**Fig. 3C**). The greatest overlap of >57,000 peptides was observed among all three strategies, while DIA: NoFAIMS had the highest number of unique peptides (>37,000), as expected. The next highest overlap was between the DIA methods, with nearly 37,000 peptides.

We calculated the correlation of fold changes between the IHCF and MCF7 cell lines in the DIA and TMT datasets at the peptide level. When comparing DIA: NoFAIMS with TMT, the correlation plot yielded a Pearson R² = 0.8438 (**Fig. 3D**). Similarly, comparing DIA: CV-35 with TMT, yielded a similar value, R² = 0.8189 (**Fig. 3E**). In general, the abundance ratios between datasets were highly correlated. To evaluate quantification quality, we assessed precision by calculating the coefficient of variation (relative standard deviation, RSD) for all replicates with no missing values. At the peptide level, we plotted the RSD distribution for all three datasets (**Fig. 3F**). The average RSDs were approximately 12%, 22%, and 10% for the DIA: NoFAIMS, DIA: CV-35, and TMT datasets, respectively. The protein-level RSD distributions reflected those at the peptide level, with values approximately half of those at the peptide level: 7%, 11%, and 5% for the DIA: NoFAIMS, DIA: CV-35, and TMT datasets, respectively (**Fig. 3G**). Overall, these findings were encouraging, demonstrating a strong correlation and high precision between the DIA and TMT methods. The results further reinforced the observation of comparable quantitative outcomes when using DIA- and TMT-based approaches.

### Similar biological findings can be extracted when using DIA and TMT data acquisition strategies

Along with the general overview of the datasets, we sought to explore the biological insights that can be teased from these three datasets. We performed principal component analysis (PCA) on the three datasets to assess global data quality (**Fig. 4A**). For all three strategies - DIA: NoFAIMS (left), DIA: CV-30 (center), and TMT (right) - we showed similar sample clustering profiles. Replicates of the four cell lines clustered together. Likewise, the clustering patterns among the three datasets were similar, with HCT116 and HeLa cells being closer and MCF7 cells being the most segregated cell line. We noted that the IHCF cells, regardless of treatment, clustered tightly together, specifically due to few measurable abundance changes at the protein level following H_2_O_2_ treatment. We also noted a similar number of differentially abundant proteins that were significantly higher (**Fig. S2A**) or lower (**Fig. S2B**) in abundance with respect to the untreated IHCF cell line. Although our current analysis revealed limited protein alterations in response to H_2_O_2_ treatment, we have obtained additional data indicating significant changes in cysteine reactivity. These findings will be explored in depth in a forthcoming study. In addition to this global profiling, we highlighted four proteins showcasing various profiles with both large and subtle changes in protein abundance.

**Figure 4:**
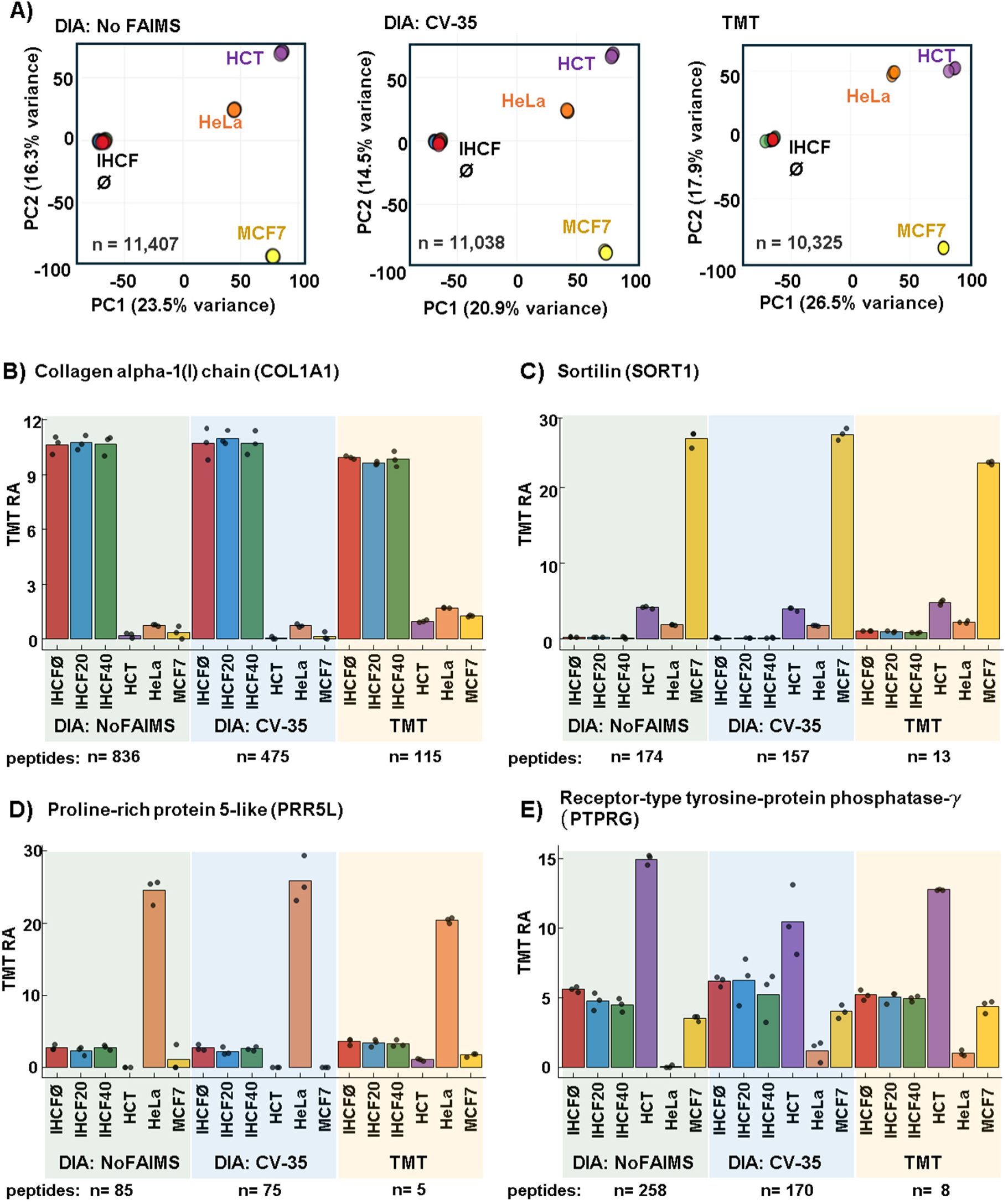
Quantitative assessment of the three data acquisition methods. Principal components analysis (PCA) plotting the first two principal components for the **A)** DIA: No FAIMS (left), DIA:CV-35 (center), and TMT (right) datasets. Example protein abundance profiles for **B)** Collagen alpha-1(I) chain (COL1A1, P02452), **C)** Sortilin (SORT1, Q99523), **D)** Proline-rich protein 5-like (PRR5L, Q6MZQ0) and **E)** Receptor-type tyrosine-protein phosphatase gamma (PTPRG, P23470).

For example, the Collagen alpha-1(I) chain (COL1A1) is highly expressed in IHCF cells and less so in the other three cell lines, which coincides with the profiles of all three datasets (**Fig. 4B**). Next, Sortilin (SORT1) is a cytokine that is a potential driver of tumorigenesis associated with cancer progression and treatment resistance [26]. Sortilin was particularly abundant in the MCF7, breast cancer cell line in all three datasets (**Fig. 4C**). This protein was present to a lesser extent in HCT116 and HeLa cell lines, however we noted that these ratios were similar in all three data sets. Next, proline-rich protein 5-like (PRR5L) is a protein that associates with the MTOR complex [27]. In all three data sets, this protein is highly abundant in HeLa cells (**Fig. 4D**). This protein, however, may be absent or of very low abundance in HCT116 and MCF7 cells, with each acquisition method showing varying degrees of background signal. In contrast, receptor-type tyrosine-protein phosphatase-g (PTPRG) showed the highest abundance in the HCT116 cell line, approximately equal abundance in IHCF and MCF7, and was very low in HeLa cells (**Fig. 4E**). PTPRG aids in maintaining stem-cell-like features and promoting oxaliplatin resistance in colorectal cancer cells [28]. We noted in these examples and throughout the datasets (such as shown in Fig. 3D and 3G), that the TMT data were expected to be impacted by ratio compression due to the inherent reporter ion interference originating from co-isolated, co-fragmenting, and co-analyzed precursors [10, 29].

### Conclusions

As quantitative mass spectrometry-based proteomics advances, the choice between TMT- and DIA-based strategies will depend on specific study requirements and available instrumentation. Both acquisition methods can provide high throughput, quantitative precision, and proteome depth through different approaches. TMT may be preferred for studies focusing on relative quantification across multiple conditions, whereas DIA may be more suitable for studies requiring comprehensive proteome coverage across many samples.

TMT’s multiplexing capability is invaluable for studies requiring relative quantification across multiple conditions, allowing simultaneous analysis of multiple samples in a single run, thus increasing throughput and efficiency. However, the method’s susceptibility to ratio compression and reporter ion interference is a notable limitation, potentially affecting quantification accuracy in complex samples. Additionally, the high cost of the labeling reagents and additional sample processing steps may impede widespread adoption. In contrast, DIA’s comprehensive data acquisition approach provides exceptional sensitivity, making it suitable for large-scale proteomic studies requiring high sensitivity. The trade-off is the increased computational complexity and processing demands required for accurate data interpretation.

Ultimately, while both strategies exhibit different strengths, they often generate comparable results. The selection of the appropriate strategy may ultimately depend on available resources, instrumentation, and the specific scientific questions to be addressed. Researchers should carefully consider these factors when designing their proteomic studies to optimize data quality and ensure relevance to their research objectives.

## ACKNOWLEDGEMENTS

We would like to acknowledge S. P. Gygi and the Taplin Mass Spectrometry Facility at Harvard Medical School for use of their mass spectrometers and software. This work was funded in part by NIH/NIGMS grant R01 GM132129 (J.A.P.) and NIH/NEI grant K99 EY035260 (X.L.). We declare no conflicts of interest.

---

**Figure S1:**
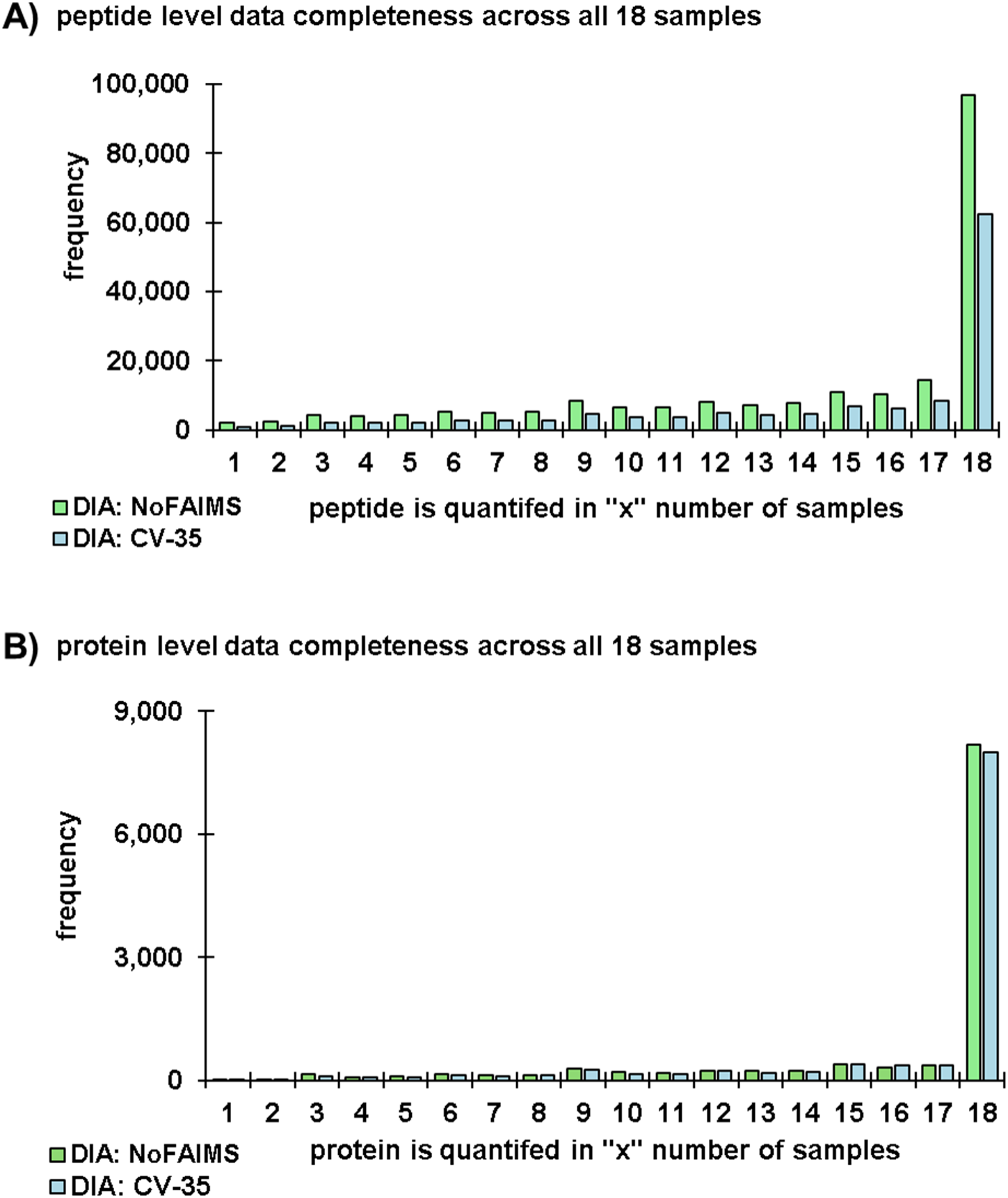
Data completion across all samples. Bar plots illustrate **A)** peptide and **B)** protein level completeness for each DIA acquisition method. These data contrasted with those in Fig. 2E and 2F, as, there, data completeness is defined by triplicate and a triplicate with no measurement in all three replicates is still considered “complete.” Here, however, a measurement for all 18 samples is required to be considered “complete.”

**Figure S2:**
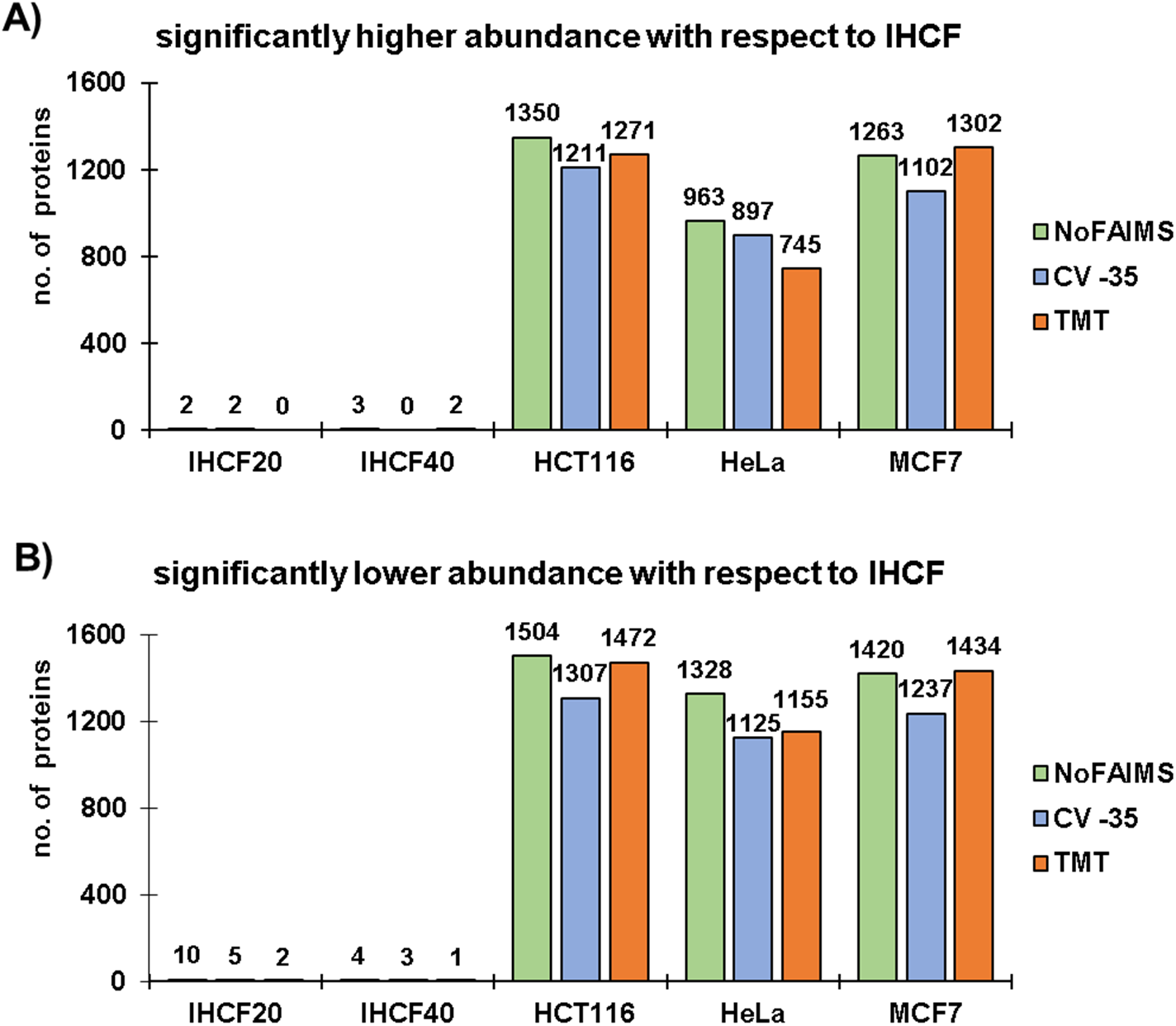
Tally of differentially abundant proteins with respect to the untreated IHCF cell line. Bar charts counting proteins that were of **A)** higher or **B)** lower abundance with statistical significance for a given condition or cell line with respect to the untreated IHCF cell line. Statistically significant differences were defined as having |log_2_(X/ IHCF)|>1 (where “X” is another cell line or treatment) and p-value <0.01.

## Notes

### Competing Interest Statement

The authors have declared no competing interest.

## REFERENCES

[1] L.C. Gillet, P. Navarro, S. Tate, H. Rost, N. Selevsek, L. Reiter, R. Bonner, R. Aebersold, Targeted data extraction of the MS/MS spectra generated by data-independent acquisition: a new concept for consistent and accurate proteome analysis, Mol Cell Proteomics 11(6) (2012) O111016717.

[2] J.D. Egertson, A. Kuehn, G.E. Merrihew, N.W. Bateman, B.X. MacLean, Y.S. Ting, J.D. Canterbury, D.M. Marsh, M. Kellmann, V. Zabrouskov, C.C. Wu, M.J. MacCoss, Multiplexed MS/MS for improved data-independent acquisition, Nat Methods 10(8) (2013) 744–6.

[3] K. Barkovits, S. Pacharra, K. Pfeiffer, S. Steinbach, M. Eisenacher, K. Marcus, J. Uszkoreit, Reproducibility, Specificity and Accuracy of Relative Quantification Using Spectral Library-based Data-independent Acquisition, Mol Cell Proteomics 19(1) (2020) 181–197.

[4] A. Thompson, J. Schafer, K. Kuhn, S. Kienle, J. Schwarz, G. Schmidt, T. Neumann, R. Johnstone, A.K. Mohammed, C. Hamon, Tandem mass tags: a novel quantification strategy for comparative analysis of complex protein mixtures by MS/MS, Anal Chem 75(8) (2003) 1895–904.

[5] P.L. Ross, Y.N. Huang, J.N. Marchese, B. Williamson, K. Parker, S. Hattan, N. Khainovski, S. Pillai, S. Dey, S. Daniels, S. Purkayastha, P. Juhasz, S. Martin, M. Bartlet-Jones, F. He, A. Jacobson, D.J. Pappin, Multiplexed protein quantitation in Saccharomyces cerevisiae using amine-reactive isobaric tagging reagents, Mol Cell Proteomics 3(12) (2004) 1154–69.

[6] D.C. Frost, T. Greer, L. Li, High-resolution enabled 12-plex DiLeu isobaric tags for quantitative proteomics, Anal Chem 87(3) (2015) 1646–54.

[7] C.S. Sauer, L. Li, Multiplexed quantitative neuropeptidomics via DiLeu isobaric tagging, Methods Enzymol 663 (2022) 235–257.

[8] N.R. Zuniga, D.C. Frost, K. Kuhn, M. Shin, R.L. Whitehouse, T.Y. Wei, Y. He, S.L. Dawson, I. Pike, R.D. Bomgarden, S.P. Gygi, J.A. Paulo, Achieving a 35-Plex Tandem Mass Tag Reagent Set through Deuterium Incorporation, J Proteome Res 23(11) (2024) 5153–5165.

[9] G.C. McAlister, E.L. Huttlin, W. Haas, L. Ting, M.P. Jedrychowski, J.C. Rogers, K. Kuhn, I. Pike, R.A. Grothe, J.D. Blethrow, S.P. Gygi, Increasing the multiplexing capacity of TMTs using reporter ion isotopologues with isobaric masses, Anal Chem 84(17) (2012) 7469–78.

[10] J.A. Paulo, J.D. O’Connell, S.P. Gygi, A Triple Knockout (TKO) Proteomics Standard for Diagnosing Ion Interference in Isobaric Labeling Experiments, J Am Soc Mass Spectrom 27(10) (2016) 1620–5.

[11] X. Liu, S.P. Gygi, J.A. Paulo, Isobaric Tag-Based Protein Profiling across Eight Human Cell Lines Using High-Field Asymmetric Ion Mobility Spectrometry and Real-Time Database Searching, Proteomics 21(1) (2021) e2000218.

[12] X. Liu, V. Rossio, J.A. Paulo, Spin column-based peptide fractionation alternatives for streamlined tandem mass tag (SL-TMT) sample processing, J Proteomics 276 (2023) 104839.

[13] T. Zhang, X. Liu, V. Rossio, S.L. Dawson, S.P. Gygi, J.A. Paulo, Enhancing Proteome Coverage by Using Strong Anion-Exchange in Tandem with Basic-pH Reversed-Phase Chromatography for Sample Multiplexing-Based Proteomics, J Proteome Res 23(8) (2024) 2870–2881.

[14] J. Navarrete-Perea, Q. Yu, S.P. Gygi, J.A. Paulo, Streamlined Tandem Mass Tag (SL-TMT) Protocol: An Efficient Strategy for Quantitative (Phospho)proteome Profiling Using Tandem Mass Tag-Synchronous Precursor Selection-MS3, J Proteome Res 17(6) (2018) 2226–2236.

[15] Y. Wang, F. Yang, M.A. Gritsenko, Y. Wang, T. Clauss, T. Liu, Y. Shen, M.E. Monroe, D. Lopez-Ferrer, T. Reno, R.J. Moore, R.L. Klemke, D.G. Camp, 2nd, R.D. Smith, Reversed-phase chromatography with multiple fraction concatenation strategy for proteome profiling of human MCF10A cells, Proteomics 11(10) (2011) 2019–26.

[16] J.A. Paulo, J.D. O’Connell, R.A. Everley, J. O’Brien, M.A. Gygi, S.P. Gygi, Quantitative mass spectrometry-based multiplexing compares the abundance of 5000 S. cerevisiae proteins across 10 carbon sources, J Proteomics 148 (2016) 85–93.

[17] S.A. Beausoleil, J. Villen, S.A. Gerber, J. Rush, S.P. Gygi, A probability-based approach for high-throughput protein phosphorylation analysis and site localization, Nature biotechnology 24(10) (2006) 1285–92.

[18] E.L. Huttlin, M.P. Jedrychowski, J.E. Elias, T. Goswami, R. Rad, S.A. Beausoleil, J. Villen, W. Haas, M.E. Sowa, S.P. Gygi, A tissue-specific atlas of mouse protein phosphorylation and expression, Cell 143(7) (2010) 1174–89.

[19] J.E. Elias, S.P. Gygi, Target-decoy search strategy for mass spectrometry-based proteomics, Methods Mol Biol 604 (2010) 55–71.

[20] J.E. Elias, S.P. Gygi, Target-decoy search strategy for increased confidence in large-scale protein identifications by mass spectrometry, Nat Methods 4(3) (2007) 207–14.

[21] G.C. McAlister, E.L. Huttlin, W. Haas, L. Ting, M.P. Jedrychowski, J.C. Rogers, K. Kuhn, I. Pike, R.A. Grothe, J.D. Blethrow, S.P. Gygi, Increasing the Multiplexing Capacity of TMTs Using Reporter Ion Isotopologues with Isobaric Masses, Analytical Chemistry 84(17) (2012) 7469–7478.

[22] J. Navarrete-Perea, X. Liu, R. Rad, J.P. Gygi, S.P. Gygi, J.A. Paulo, Assessing interference in isobaric tag-based sample multiplexing using an 18-plex interference standard, Proteomics 22(7) (2022) e2100317.

[23] D.K. Schweppe, S. Prasad, M.W. Belford, J. Navarrete-Perea, D.J. Bailey, R. Huguet, M.P. Jedrychowski, R. Rad, G. McAlister, S.E. Abbatiello, E.R. Woulters, V. Zabrouskov, J.J. Dunyach, J.A. Paulo, S.P. Gygi, Characterization and Optimization of Multiplexed Quantitative Analyses Using High-Field Asymmetric-Waveform Ion Mobility Mass Spectrometry, Anal Chem 91(6) (2019) 4010–4016.

[24] J. Saba, E. Bonneil, C. Pomies, K. Eng, P. Thibault, Enhanced sensitivity in proteomics experiments using FAIMS coupled with a hybrid linear ion trap/Orbitrap mass spectrometer, J Proteome Res 8(7) (2009) 3355–66.

[25] K.E. Swearingen, M.R. Hoopmann, R.S. Johnson, R.A. Saleem, J.D. Aitchison, R.L. Moritz, Nanospray FAIMS fractionation provides significant increases in proteome coverage of unfractionated complex protein digests, Mol Cell Proteomics 11(4) (2012) M111014985.

[26] S. Rhost, E. Hughes, H. Harrison, S. Rafnsdottir, H. Jacobsson, P. Gregersson, Y. Magnusson, P. Fitzpatrick, D. Andersson, K. Berger, A. Stahlberg, G. Landberg, Sortilin inhibition limits secretion-induced progranulin-dependent breast cancer progression and cancer stem cell expansion, Breast Cancer Res 20(1) (2018) 137.

[27] D.G. Ozturk, M. Kocak, A. Akcay, K. Kinoglu, E. Kara, Y. Buyuk, H. Kazan, D. Gozuacik, MITF-MIR211 axis is a novel autophagy amplifier system during cellular stress, Autophagy 15(3) (2019) 375–390.

[28] C. Fu, X. Li, L. Han, M. Xie, S. Ouyang, LncRNA PTPRG-AS1 maintains stem-cell-like features and promotes oxaliplatin resistance in colorectal cancer via regulating the miR-665 and STAT3 axis, Molecular & Cellular Toxicology 20(1) (2024) 35–45.

[29] J.P. Gygi, Q. Yu, J. Navarrete-Perea, R. Rad, S.P. Gygi, J.A. Paulo, Web-Based Search Tool for Visualizing Instrument Performance Using the Triple Knockout (TKO) Proteome Standard, J Proteome Res 18(2) (2019) 687–693.

